# metagWGS, a comprehensive workflow to analyze metagenomic data using Illumina or PacBio HiFi reads

**DOI:** 10.1101/2024.09.13.612854

**Authors:** Jean Mainguy, Maïna Vienne, Joanna Fourquet, Vincent Darbot, Céline Noirot, Adrien Castinel, Sylvie Combes, Christine Gaspin, Denis Milan, Cécile Donnadieu, Carole Iampietro, Olivier Bouchez, Géraldine Pascal, Claire Hoede

## Abstract

**Background:** To study communities of micro-organisms taxonomically and functionally, metagenomic analyses are now often used. If there is no reference gene catalogue, a *de novo* approach is required. Because genomes are easier to interpret than contigs, the recovery of metagenome-assembled genomes (MAGs) by binning of contigs from metagenomic data has recently become a common task for microbial studies. However, during this process, there is a significant loss of information between the assembly and the binning of contigs. This is why it is important to produce taxonomic and functional matrices for all contigs and not just those included in correct bins. In addition, Pacbio HiFi reads (long and of good quality) are now a possible, albeit more expensive, alternative to short Illumina reads. We therefore developed a workflow that is easy to install with dependencies fixed using singularity images and easy to use on a computing cluster, that is capable of analyzing either short or long reads, and that should allow analysis at the contig and/or bin level, depending on the user’s choice.

Following is a presentation of metagWGS, a fully automated workflow for metagenomic data analysis. It uses a new tool for refining bins (called Binette) that we will demonstrate is more efficient than competing tools.

**Methods:** metagWGS is a Nextflow workflow distributed with two singularity images and complete documentation to facilitate its installation and use.

Because the main original features of metagWGS concern binning (short and long reads) and the analysis of HiFi reads, we compared metagWGS with the MAG construction workflow proposed by PacBio to a public dataset used by Pacbio to promote its workflow.

**Results:** metagWGS differs from existing workflows by (i) offering flexible approaches for the assembly; (ii) supporting short reads (Illumina) or PacBio HiFi reads; (iii) combining multiple binning algorithms with a new bin refinement tool, referred to as “Binette”, to achieve high-quality genome bins; and (iv) providing taxonomic and functional annotation for all genes, all contigs built and bins.

metagWGS produces more medium (708) and high-quality (255) bins on 11 public metagenomic samples from human gut data than the Pacbio HiFi dedicated workflow, referred to as the HiFi-MAGS-pipeline (659 medium quality bins and 231 high quality bins), primarily due to the better performance of Binette.

## Introduction

Metagenomic analyses provide access to the taxonomic and functional diversity of the communities studied. In addition, depending on the dataset, it is possible to reconstitute the genomes of the species that are easiest to assemble. A number of approaches exist, including those dealing solely with reads, those focusing on contigs and those analyzing MAGs alone. For the most extensively studied environments, an approach using a homology search on a reference catalogue is effective, but when little is known about the ecosystem, it is worth implementing a *de novo* strategy.

For our analyses of as yet little-studied communities, we chose to obtain as much information as possible, which meant first constructing taxonomic and functional abundance matrices from as many contigs as possible, then trying to group the contigs that could be grouped into bins, and subsequently obtaining the MAGs (metagenome-assembled genomes) and their abundance in each sample. We also wanted to be able to install and run a single tool for all the necessary steps, i.e., cleaning, checking data quality, assembly, mapping, taxonomic and functional annotation of contigs, quantification of their abundance, construction of the functional abundance matrix (the catalogue of genes with their abundance in the samples), binning, dereplication of the bins and their annotation and, finally, building the MAG abundance matrices.

Current tools are designed to reconstruct and analyze MAGs and do not output taxonomic and functional profiles for all the contigs (MAG (*Krakau et al., 2022*), MetaWRAP (*Uritskiy, DiRuggiero & Taylor, 2018*), Veba (*Espinoza & Dupont, 2022*), Atlas (*Kieser et al., 2020*)). Anvi’o is capable of generating these profiles but is designed to allow interactive work in a Web browser, and the necessary tools are not integrated in a pre-configured workflow for launching the entire analysis in one command on a computing cluster.

That is why we present metagWGS, a workflow implemented in Nextflow DSL2 (*Di Tommaso et al., 2017*) that is able to analyze whole shotgun sequence metagenomic data. Its main original features are: (i) it is able to deal with Illumina short reads or PacBio HiFi reads; (ii) it is comprehensive as it analyzes contigs, genes and MAGs (metagenome-assembled genomes). In particular it produces a taxonomic abundance table from the contigs and from the MAGs. A list of non-binned contigs is provided; (iii) it also produces a functional abundance table from the catalogue of genes found in the contigs; (iv) it includes an improved algorithm for automatic bin refinement, Binette (J. Mainguy, C. Hoede, 2024) available here: https://github.com/genotoul-bioinfo/Binette. metagWGS is publicly-available at: https://forgemia.inra.fr/genotoul-bioinfo/metagwgs with complete documentation. The pipeline strikes a balance between providing flexibility for users to customize their analyses (e.g., choice of parameters, steps to be executed) and maintaining accessibility for the user, thereby capable of addressing a broad spectrum of biological questions in shotgun metagenomic studies. We also provide user support and plan to offer training courses.

Because the main original features of metagWGS concern binning (short and long reads) and the analysis of HiFi reads, we compared metagWGS with the MAG construction workflow proposed by PacBio, referred to as the HiFi-MAGS-pipeline, using a public dataset composed of 11 public metagenomic samples from human gut.

## Materials and methods

### Datasets

We compared binning step results between metagWGS and the HiFi-MAGS-pipeline provided by PacBio (https://github.com/PacificBiosciences/pb-metagenomics-tools/blob/master/docs/Tutorial-HiFi-MAG-Pipeline.md) using 11 public metagenomic samples from human gut, a publicly-available HiFi dataset from the ncbi project ID: PRJNA754443 (Run IDs: SRR15489010, SRR15489009, SRR15489020, SRR15489019, SRR15489018, SRR15489017, SRR15489016, SRR15489015, SRR15489014, SRR15489013 and SRR15489011). The objectives of the project that generated these data were to evaluate short-read and long-read sequencing approaches for metabarcoding or whole genome metagenomic analysis to maximize the utility of clinical microbiome data (*Gehrig et al., 2022*). We used only shotgun data for our comparison.

We compared the binning refinement tools Binette, DAS Tool, and the metaWRAP bin refinement module using the simulated mouse gut metagenome data (https://doi.org/10.4126/FRL01-006421672), which was released in preparation for the second round of the CAMI II challenges (*Meyer et al., 2022*). This challenge allows the evaluation of metagenomic tools on realistic and complex datasets with long- and short-read sequences, created computationally.

### Description of the workflow

metagWGS is split into eight different steps that correspond to different parts of the bioinformatics analysis. Many of these steps are optional and their necessity depends on the desired analysis.

Step 1 is the cleaning step that trims adapter sequences and deletes low-quality reads with Cutadapt v4.2 (*Martin, 2011*) and Sickle v1.33 (Joshi et al., 2011), suppresses host contaminants with BWA-MEM2 v2.2.1 (*Vasimuddin et al., 2019*) or Minimap2 v2.24 (*Li, 2018*) and Samtools v1.15.1 (*Danecek et al., 2021*), controls the quality of raw and cleaned data (FastQC v0.11.9), and makes a taxonomic classification of cleaned reads with Kaiju MEM v1.9.2 (*Menzel et al., 2016*), kronaTools v2.8.1 (*Ondov et al., 2011*), plot_kaiju_stat.py and merge_kaiju_results.py. plot_kaiju_stat.py and merge_kaiju_results.py are in-house python scripts. Plot_kaiju_stat.py produces an html page with the match length distribution plots and merge_kaiju_results.py merge kaiju results by taxon level for all samples. The aim of this analysis of reads by kaiju is to highlight any contamination of the data by rapid analysis. By default, the database used is refseq_bacteria to make it computationally reasonable.

Step 2 is the assembly step. Short reads can be assembled by metaSpades v3.15.5 (*Prjibelski et al., 2020*) or megahit v1.2.9 (*Li et al., 2015*; *Li et al., 2016*). The default is metaSpades. HiFi reads can be assembled by Hifiasm-meta v0.3 (*Feng et al., 2022*) or metaFlye v2.9.1 (*Kolmogorov et al., 2020*). The default is Hifiasm-meta. Users can choose between them in the configuration file (nextflow.config) or via a workflow option. Short reads are aligned against contigs with BWA-MEM2, and duplicated reads are removed by samtools. HiFi reads are aligned by Minimap2. In both cases, samtools is used to compute alignment statistics and coverage for each nucleotide of contigs. By filling in the “group” field in the sample sheet given as the workflow input, it is possible to request co-assembly of all or just some of the samples. We recommend using megahit to co-assemble a large quantity of samples in short reads. If the assembly is complicated and requires, for example, *in silico* normalization of the reads, metagWGS can take assemblies previously made outside the workflow as input.

Step 3 allows the assembly to be filtered on the normalized (by library size) count of the reads mapped on the contigs by an in-house script: Filter_contig_per_cpm.py. The threshold for minimal counts per millions of reads can be configured by a dedicated parameter. This step can be skipped. In this step, the final assembly quality is evaluated by metaQUAST v5.2.0 (*Mikheenko et al., 2016*).

Step 4 consists of the structural annotation of the contigs, CDS are annotated by Prodigal v2.6.3, rRNA by Barrnap v0.9, tRNA by tRNAscan-SE v2.0.11 (*Chan et al., 2021*). The three annotations are merged in one GFF file per sample by an in-house script: merge_annotations.py.

Step 5 prepares taxonomic annotation of genes by aligning the protein sequence of genes against a protein database with DIAMOND v2.0.15 (*Buchfink, Reuter & Drost, 2021*). The recommended protein bank is nr from ncbi.

Step 6 makes a sample and global clustering of genes with cd-hit-est v4.8.1 (*Li & Godzik, 2006*; *Fu et al., 2012*), and an in-house script (cd_hit_produce_table_clstr.py) produces a table of the genes making up each cluster. Reads mapped on annotations are quantified by featureCounts and an in-house script, referred to as Quantification_clusters.py, adds up the number of reads of the genes making up each cluster and writes a file containing a correspondence table between global cluster id and gene id. Genes are functionally annotated by eggNOG-mapper v2.1.9 (*Cantalapiedra et al., 2021*; *Huerta-Cepas et al., 2019*) and two in-house scripts: merge_abundance_and_functional_annotations.py and quantification_by_functional_annotation.py compile functional annotation of each representative cluster, quantification for each sample of all genes in each cluster and the best DIAMOND hit.

Step 7 taxonomically affiliates the genes by using DIAMOND against protein database results (nr by default) and a LCA (Low Common Ancestor) algorithm implemented in aln2taxaffi.py (in-house script). In this script, we parse DIAMOND output and keep hits with the best score for each gene (query) (score = query_coverage/100 x percent_identity/100) if and only if query_coverage >= 70% and percent_identity >= 60%. We retrieve taxon ID for each accession number corresponding to best hits. For each best hit (it is possible to have multiple best hits), we associated a weight by taxonomy ranks. weight = (score - RANKS_TO_MIN_SCORE[rank]) / (1.0 - RANKS_TO_MIN_SCORE[rank]); weight = max(weight, 0.0). RANKS_TO_MIN_SCORE = {’superkingdom’: 0.4, ‘phylum’: 0.5, ‘class’: 0.6, ‘order’: 0.7, ‘family’: 0.8, ‘genus’: 0.9, ‘species’: 0.95}. For each taxonomy rank, the script sums weights and computes the best_taxid_score/sum of weight; if the result is > 0.9, we keep this taxonomy consensus; otherwise, we write that we were unable to find a taxonomy consensus for this gene in the gene taxonomic affiliation file. We use the same algorithm to find the consensus taxonomy for each contig based on the best DIAMOND hits of the genes annotated on the contigs. We provide a file with the taxonomy at each rank and the associated score to help users to understand the final choice. By using the mapping results, we quantify taxonomic abundance, write a matrix of relative abundance for each sample and provide various plots like krona and histogram of relative abundance for each taxonomic level.

Step 8 is the final step. Users can choose between three mapping strategies to compute co-abundances for the binning of contigs (Individual, group or all). Individual: only reads of the sample are mapped on corresponding assembly; group: reads of a sample group (defined in sample sheet) are mapped against all assemblies of this group; all: all reads from all samples are mapped against all assemblies. BWA-MEM2 is used to map short reads, minimap2 for HiFi reads. Binning is done using three tools: MetaBat2 v2.15 (*Kang et al., 2019*), MaxBin2 v2.2.7 (*Wu et al., 2016*) and CONCOCT v1.1.0 (*Alneberg et al., 2014*) and refined by Binette v1.0.1 (*Mainguy & Hoede, 2024*). For HiFi reads, we also use the circular contigs as bins. If several bins have at least one contig in common, Binette generates new hybrid bins based on the intersection, the difference and union of bins to expand the range of possible bins, and then compares the quality of each hybrid bin by checkM2 v1.0.1 (Chklovski, Parks, Woodcroft., 2023) to choose the best one. Bins obtained for each sample are then dereplicated by dRep v3.0.0 (*Olm et al. 2017*), and MAGs (metagenome-assembled genomes) are evaluated by checkM2. The resulting bins are taxonomically affiliated by GTDB-tk v2.1 (*Chaumeil et al., 2022*). Finally we compute and provide a matrix of abundance for bins.

All steps are launched one after another by default. The user can choose to stop at a step or skip a step with dedicated parameters.

Finally, a multiQC v1.14 (*Ewels et al., 2016*) report is produced that brings together numerous statistics and graphs from the different steps. A file with the versions of the databases and software used is also provided to the user.

### Implementation and reproducibility

We used Nextflow DSL 2 to allow job management and error recovery. metagWGS comes with two singularity images that contain all the dependencies. We provide sources and complete documentation through the *GitLab* repository (https://forgemia.inra.fr/genotoul-bioinfo/metagwgs). We use and provide functional tests in the *GitLab* repository (https://forgemia.inra.fr/genotoul-bioinfo/metagwgs-test-datasets), input and expected output datasets. MetagWGS is developed with python 3.10.8.

### Comparative analysis of binning results from HiFi-MAGS-Pipeline and metagWGS

To make this comparison, we ran version 2.4.2 of metagWGS (doi: 10.5281/zenodo.10007876) without filtering human reads, with hifiasm v0.13-r308 as the assembler, without filtering contigs on their reads depth and with cross-mapping for the binning. We then retrieved the assemblies produced for each sample and used them as input to the PacBio HiFi-MAGS-pipeline 2.0.2. We used default parameters for the HiFi-MAGS-pipeline except for--environment=human_gut and bin quality filters min_completeness: 50, max_contamination: 10, max_contigs: 200, and the same GTDB is used for GTDB-tk for both analyses in order to be able to compare the number of bins produced between the two workflows. Both workflows were launched on a cluster (with Slurm as the scheduler) whose nodes have the following specifications: Bullx R424-E4 processor: Intel(R) Xeon(R) CPU E5-2683 v4 @ 2.10GHz; 64 threads by node; 512GB RAM by node; with Linux 64 bits based on CentOS 7 distribution and Interconnexion Infiniband (FDR); the number of cores used depends on the configuration files present in the workflow source.

### Binning benchmark on simulated dataset

We compared Binette v1.0.0 with two other bin refinement tools, metaWRAP bin refinement tool v1.3.1 and DAS Tool v1.1.7 using the simulated mouse gut metagenome data released in preparation for the second round of CAMI II challenges (*Meyer et al., 2022*). To make the comparison, we followed the step included in this nature protocol describing the binning benchmark (*Meyer et al., 2021*).

As input of the bin refinement tools, we used the results produced from the cross-sample gold-standard assembly with the binners MaxBin v.2.2.7, MetaBAT v.2.12.1 and CONCOCT v.1.0.0 available in the CAMI tool result repositories on Zenodo (https://zenodo.org/communities/cami). The three bin refinement tools were run using ten CPUs on the same node of the computing cluster described above.

Binning results from the individual binners (MaxBin 2, MetaBAT 2 and CONCOCT) and refinement tools (Binette, metaWRAP and DAS Tool) were evaluated using AMBER v2.0.4 (*Meyer et al., 2018*), which generated binning quality metrics based on the ground truth of the simulated data. Main metrics are purity, completeness and the F1 score (the F1 score summarizes average bin purity and genome completeness).

## Results

### Pipeline overview

metagWGS is a highly modular workflow consisting of eight steps (Fig. 1). It is possible to stop at the desired steps or skip certain steps. Some tool calls are also optional inside each step.

**Fig. 1:**
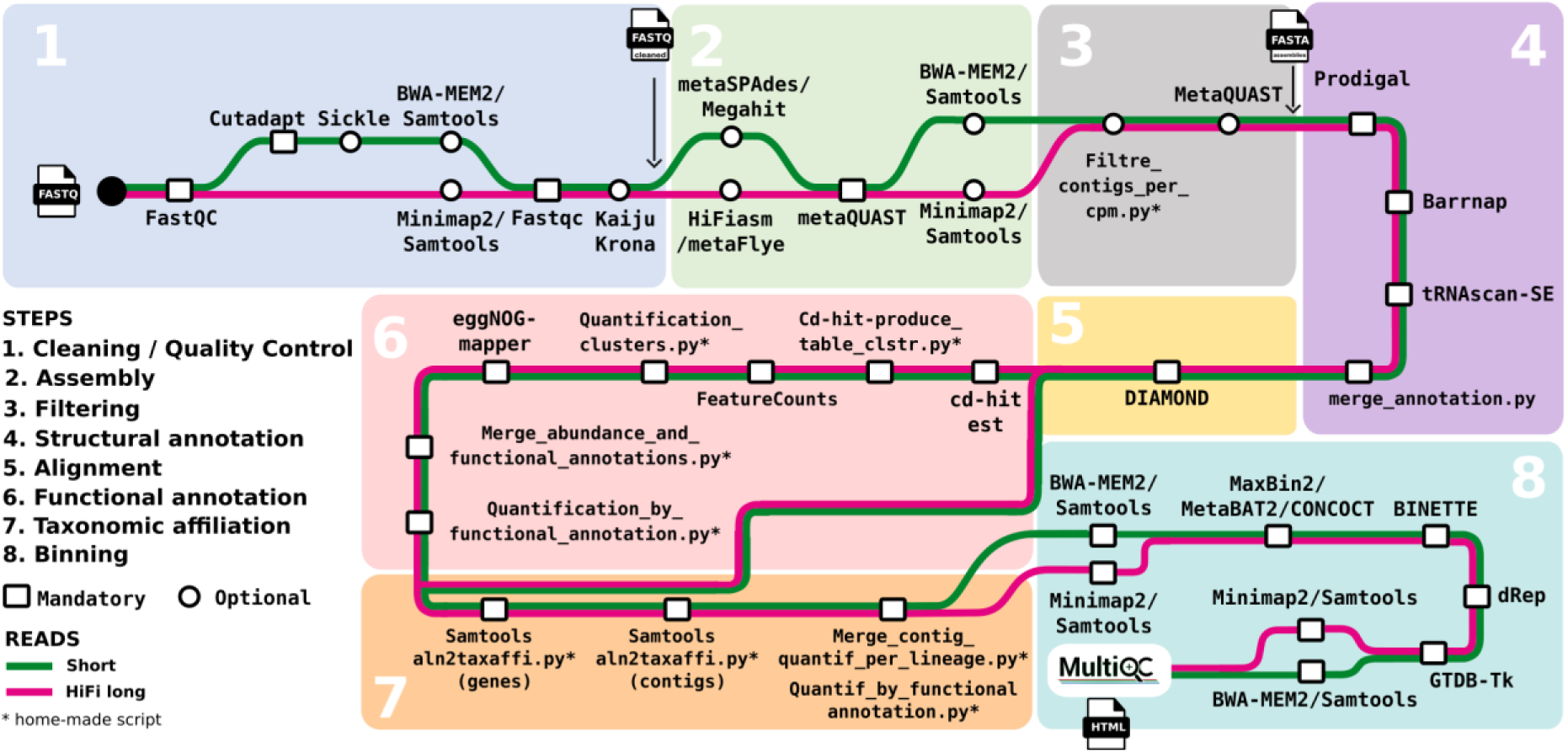
Workflow summary diagram. The green path corresponds to the short reads, the magenta one to the HiFi reads.

To use metagWGS, it is necessary to provide the metagenomic whole genome shotgun data you want to analyze (Illumina HiSeq3000, NovaSeq sequencing, 2*150bp or PacBio HiFi reads). If it is necessary to assemble data outside metagWGS, you can also provide the assembly as input. Input files and groups (if you want to perform co-assembly or cross alignment for binning) are specified in a sample sheet. All parameters are described in the usage documentation available in the gitlab repository of the workflow. You can also specify a path to databases or all other parameters in config files. We provide files “nextflow.config” and “genotoul.config” for you to work from in the gitlab repository. The “base.config” file also available in the repository provides a list of typical resource reservations on which to base a standard metagenomics project on our cluster for all steps of the workflow.

metagWGS generates a large number of outputs organized by stage and described in detail in the output documentation provided in the gitlab repository.

At the end of the workflow, it provides an html report from MultiQC in which you will find some interesting plots like, for example, Supplementary Fig. 1 that highlights the contig size for each sample, Supplementary Fig. 2 that represents bins quality for each sample, and Supplementary Fig. 3 showing the quality of dereplicated bins in terms of completeness and contamination.

### Comparison with other popular metagenomic WF

We developed metagWGS because very few workflows (Anvi’o only) make it possible to follow all levels of analysis (reads, contigs, functions and MAGs). By enabling users to produce taxonomic and functional matrices for all contigs, our workflow allows for thorough verification at each analysis step independently. In particular, several workflows are unable to output the functions carried by all the contigs, particularly those that are not binned, or their taxonomic affiliation (see Supplementary Table 1 for details). MAG (*Krakau et al., 2022*) from nf-core produces and annotates MAGs but does not produce contig outputs. In addition, it does not annotate rRNAs or tRNAs, which are measures of metagenomic assembly quality for the ENA (European Nucleotide Archive). metaWRAP is no longer maintained and like the MAG pipeline, it focuses on bins and does not annotate contigs that do not belong to any bin. The HiFi-MAGS-pipeline is dedicated to HiFi reads and performs only the binning step. VEBA (*Espinoza & Dupont, 2022*) is interesting because it can analyze eukaryotic, prokaryotic and viral genomes, whereas metagWGS is optimized for bacteria. However, according to the documentation, VEBA does not annotate taxonomically or functionally non-binned contigs either.

Atlas (*Kieser et al., 2020*) produces the functional gene catalogues from all the contigs and the MAG, with taxonomic affiliation, functional annotation and quantification. However, it seems that it does not make the taxonomic affiliation of all the contigs but only of the bins.

The Anvi’o software ecosystem (Eren, Kiefl & Shaiber, 2021) is designed to allow interactive work in a Web browser. As a result, it may be difficult to run certain resource-intensive steps on a laptop or, on the contrary, to run the visualization part on a computing cluster. Moreover, there is no pre-configured workflow for launching the entire analysis in sequence, from reads to MAGs, via contigs and gene catalogues, even if the tools are all there individually. However, Anvi’o is very useful for manually refining bins via its graphical interface.

Finally, while MAG and Atlas can perform hybrid assembly with both short reads and long reads from Oxford Nanopore Technologies, they cannot natively assemble PacBio HiFi reads. The only workflow that can natively assemble PacBio HiFi reads is metagWGS (Supplementary Table 1).

### Using metagWGS on HiFi reads; comparison with HiFi-MAGS-pipeline results

We used 11 public metagenomic samples from human gut data sequenced with PacBio Sequel II (HiFi reads) as input to metagWGS 2.4.2 (accession numbers are given in the Material and Methods section).

We obtained very good assemblies with N50 between 92.2 Kpb and 529.8 Kpb. Assembly length ranged between 142.8 Mpb and 442.7 Mpb, and the percentage of mapped reads on the assemblies for each sample was between 95.2% and 97.8% (see Supplementary Table 2 for details and Supplementary Fig. 1 to visualize the distribution of contig sizes from the metagWGS’s multiQC report).

We used these assemblies as input for the PacBio HiFi-MAGS-pipeline 2.0.2 and compared the results with metagWGS 2.4.2 results concerning MAGs.

The first thing we observed was that the default resource reservation settings (that, unfortunately, users use as their first option) for the HiFi-MAGS-pipeline are too large. For example, for semibin2 analysis, we used only 2.80% of the 96GB of allocated memory. To reduce waiting times and to avoid penalizing other users, it is therefore crucial to reserve resources as close as possible to requirements by configuring the resources in the workflow configuration file.

On the dataset used here and with default resource reservation settings, metagWGS used 4307 CPU hours (the equivalent of 179 days), and lasted 3.5 days in user time for the entire workflow from step 1 (cleaning reads) to step 8 (binning) on our computing infrastructure. The binning step alone took 24 hours and 17 minutes in user time (and around 415 CPU hours). For the binning alone, the PacBio workflow also took 24 hours on the same infrastructure in user time with default resource reservation, but 44 hours in CPU time.

The HiFi-MAGS-pipeline then produces a set of bins for each sample. On the contrary, metagWGS uses dRep (*Olm et al. 2017*) to produce a final set of bins with the best bins when they are shared by several samples at the “species” level (95% Average Nucleotide Identity (ANI) by default; this value can be changed).

In our experience, nearly 50% of contigs are not used in any bins, even when the assembly is of good quality. For example, for these high-quality assemblies, we obtain between 63.5% and 34.3% of the cumulative length of assemblies contained in unbinned contigs (see Supplementary Fig. 2). That is why metagWGS lists unbinned contigs, unlike the HiFi-MAGS-pipeline. metagWGS builds more medium-quality bins with completeness > 50% and contamination < 10% than the HiFi-MAGS-pipeline for eight of the 11 samples. For all samples, MetagWGS builds as many or more high-quality bins (completeness > 90% and contamination < 5%) than the HiFi-MAGS-pipeline (see Table 1).

**Table 1:** Number of bins produced by metaGWGS and the HiFi-MAGS-pipeline for 11 public samples from a publicly available HiFi dataset. Number of medium-quality bins for each sample (with more than 50% completeness and less than 10% contamination) by the two workflows and number of high-quality bins produced for each sample with more than 90% completeness and less than 5% contamination by the two workflows. These numbers are produced from the output of each of the two software packages, and are therefore obtained with checkM2. For each bin qualification, the highest number is colored in yellow; the lowest is dark purple. When the two numbers are the same, values are colored in light purple.

|  | Mediumquality bins |  | High-quality bins |  |
| --- | --- | --- | --- | --- |
| sample | metagWGS | HiFi-MAGS-pipeline | metagWGS | HiFi-MAGS-pipeline |
| humanGut_1<br>(SRR15489020) | 46 | 50 | 20 | 20 |
| humanGut_2<br>(SRR15489019) | 53 | 53 | 19 | 17 |
| humanGut_3<br>(SRR15489018) | 105 | 93 | 45 | 43 |
| humanGut_4<br>(SRR15489017) | 90 | 89 | 25 | 19 |
| humanGut_5<br>(SRR15489016) | 30 | 28 | 6 | 5 |
| humanGut_6<br>(SRR15489015) | 65 | 50 | 21 | 17 |
| humanGut_7<br>(SRR15489014) | 38 | 34 | 18 | 15 |
| humanGut_8<br>(SRR15489013) | 43 | 38 | 15 | 14 |
| humanGut_9<br>(SRR15489011) | 79 | 68 | 24 | 23 |
| humanGut_10<br>(SRR15489010) | 84 | 80 | 29 | 25 |
| humanGut_11<br>(SRR15489009) | 75 | 76 | 33 | 33 |
| **Total** | **708** | **659** | **255** | **231** |

|  | Medium quality |  | High quality |  |
| --- | --- | --- | --- | --- |
| sample | metagWGS | HiFi-MAGS-pipeline | metagWGS | HiFi-MAGS-pipeline |
| humanGut_1<br>(SRR15489020) | 46 | 50 | 20 | 20 |
| humanGut_2<br>(SRR15489019) | 53 | 53 | 19 | 17 |
| humanGut_3<br>(SRR15489018) | 105 | 93 | 45 | 43 |
| humanGut_4<br>(SRR15489017) | 90 | 89 | 25 | 19 |
| humanGut_5<br>(SRR15489016) | 30 | 28 | 6 | 5 |
| humanGut_6<br>(SRR15489015) | 65 | 50 | 21 | 17 |
| humanGut_7<br>(SRR15489014) | 38 | 34 | 18 | 15 |
| humanGut_8<br>(SRR15489013) | 43 | 38 | 15 | 14 |
| humanGut_9<br>(SRR15489011) | 79 | 68 | 24 | 23 |
| humanGut_10<br>(SRR15489010) | 84 | 80 | 29 | 25 |
| humanGut_11<br>(SRR15489009) | 75 | 76 | 33 | 33 |
| **Total** | **708** | **659** | **255** | **231** |

From all these 708 bins, metagWGS obtains 246 MAGs dereplicated by dRep to 95% ANI (see Supplementary Table 3). The HiFi-MAGS-pipeline does not dereplicate bins to build a unique set of MAGs for all samples.

Since Binette uses checkM2 to optimize bin construction, it is preferable to use another tool to assess the quality of the output bins. We therefore produced the same table as Table 1, using checkM1 (*Parks et al., 2014*) to evaluate the quality of the bins obtained by each of the two workflows (see Supplementary Table 4). The difference is smaller, but still to the advantage of metagWGS (263 high-quality bins vs. 240).

### Benchmark of Binette with DAS Tool and metaWRAP bin refinement tool

To further investigate the binning step, we compared three tools: (i) Binette v1.0.0; (ii) the metaWRAP bin refinement tool v1.3.1; and (iii) DAS Tool v1.1.7. These comparisons were based on the results from three individual binners: MaxBin v2.2.7, MetaBAT v2.12.1, and CONCOCT v1.0.0, using simulated mouse gut metagenome data from the second round of the CAMI II challenge.

Although CONCOCT has the highest average completeness (84.7%), this advantage is offset by very low average purity (59.5%) (Supplementary Table 5). Metabat2, on the other hand, has the lowest average completeness (56.1) and the lowest F1 score (0.694). Maxbin2 has the best accuracy at the sample level (73.3) but has intermediate value for all the other metrics. metaWRAP has the best purity for this dataset (99.3%) but also has the second lowest completeness (58.3%). The DAS Tool performs well on all metrics but is not the best on any of them. Binette, on the other hand, has the best F1 score (0.789) (Supplementary Table 5).

In the context of recovering high-quality genomes (completeness > 90% and contamination < 10%), metaWRAP and Binette outperformed the DAS Tool, recovering 361 and 382, respectively, compared to DAS Tool’s 354 high-quality genomes. Notably, individual binning tools (CONCOCT, Metabat 2 and Maxbin 2 had a lower ability to recover high-quality genomes compared to the refinement tools (DAS Tool, Binette and MetaWRAP) (Fig. 2).

**Fig. 2:**
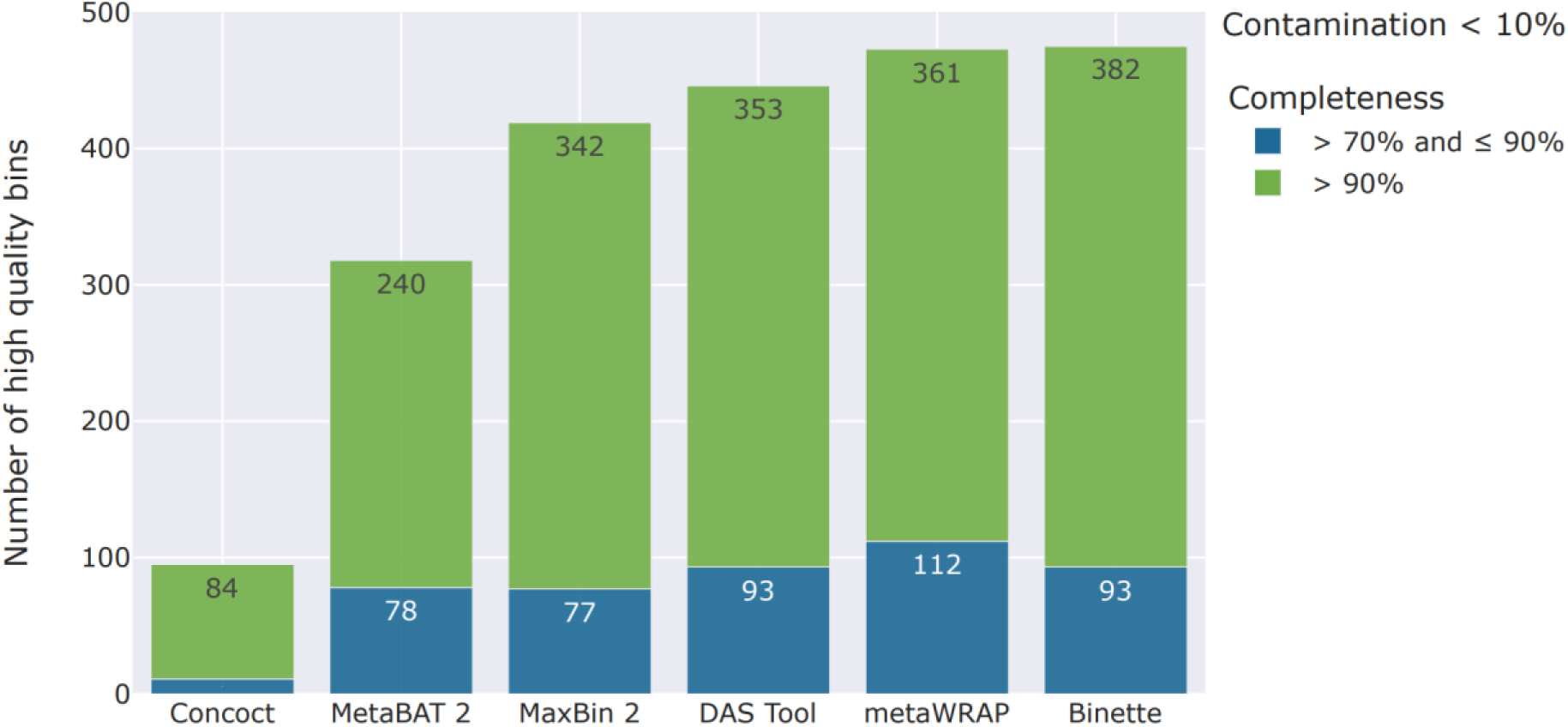
Evaluation of different binning strategies on simulated mouse gut metagenome data from the CAMI II challenges. The bar plot displays the count of high-quality bins with contamination < 10% and completeness > 70% and > 90%. DAS Tool, metaWRAP and Binette results were generated based on bins from Concoct, MetaBAT 2, and MaxBin 2.

Binette (*Mainguy & Hoede, 2024*) is an improved implementation of metaWRAP (*Uritskiy, DiRuggiero & Taylor, 2018*) refinement bin steps. From the input bin sets, metaWRAP generates new hybrid bins based solely on the intersection of bins. In contrast, Binette takes a more comprehensive approach, incorporating the difference and union of bins to expand the range of possible bins (see Fig. 1 of *Mainguy & Hoede, 2024*).

Binette and metaWRAP show comparable performance in recovering high-quality genomes for this dataset, but the average completeness of bins for metaWRAP is the second lowest in our comparison (58.3%), whereas it is 68.6% for Binette. Indeed, Binette offers a good compromise summarized by the F1 score (0.789) and the 382 high-quality genomes produced. In terms of execution time, Binette (3 hours and 7 minutes) is about 17 times faster than metaWRAP (2 days, 3 hours and 28 minutes) on this dataset. In comparison, the DAS Tool, which is slightly faster than Binette (1 hour and 10 minutes), also has a good F1 score of 0.771. However, it produces fewer high-quality genomes (353) than Binette (382).

## Discussion

### Computation time

Although the HiFi-MAGS-pipeline outperformed metagWGS in two samples, overall metagWGS succeeded in constructing 708 medium-quality MAGs on this dataset compared with 659 for the HiFi-MAGS-pipeline, i.e., 7% more.

metagWGS uses cross-alignment (i.e., it aligns the reads of all the samples on all the assemblies), which significantly improves binning but consumes CPU time. Consequently, on the dataset used here, metagWGS used 4307 CPU hours (the equivalent of 179 days), but lasted 3.5 days in user time for the entire workflow from step 1 to step 8 on our computing infrastructure thanks to our parallelization and resource reservation strategy (by default throughout the workflow). The binning step alone took 24 hours and 17 minutes in user time (and around 415 CPU hours).

For the binning alone, the PacBio workflow also took 24 hours on the same infrastructure in user time with default resource reservation, but 44 hours in CPU time. The ratio of CPU time to user time could certainly be improved in the Hifi-MAG-pipeline by improving parallelization and adapting the quantity of reserved resources.

MetagWGS uses three binning tools (MetaBAT 2, MaxBin 2 and Concoct), whereas the Pacbio pipeline uses only two (metabat2 and semibin2). Using several binning tools means you can benefit from the advantages of each of them. Further testing of combinations of binning tools to improve performance without increasing resource consumption too much would be interesting.

### Comparison of Binette with the DAS Tool and metaWRAP bin refinement tool

The Hifi-MAG-pipeline uses the DAS Tool (*Sieber et al., 2018*) to refine bins, whereas metagWGS uses Binette. The main difference between those refinement tools is that the DAS Tool, unlike Binette, does not build new hybrid bins between bins with common contigs before calculating their quality score. This probably explains why the Hifi-MAG-pipeline produces less high- and medium-quality bins than Binette. Indeed, Binette expands the range of possible bins by building hybrid bins.

We have compared bin refinement tools and the bin sets used as inputs to highlight the added value of bin refinement tools, and compared Binette, DAS Tool and MetaWRAP. Notably, we have shown that individual binning tools had a lower ability to recover high-quality genomes compared to the refinement tools (Fig. 2). These results highlight the importance of using refinement tools to improve overall binning results.

Concerning the comparison between the three bin refinement tools, Binette offers a good compromise summarized by the best F1 score (0.789) and the 382 high-quality genomes produced. MetaWRAP shows relatively good performance in recovering high-quality genomes, has the best average purity, but its average completeness of bins is the second lowest for this dataset. In terms of execution time, Binette was about 17 times faster than metaWRAP. This huge difference can be attributed to Binette’s use of CheckM2, which is faster than CheckM1 (as demonstrated in Chlovski et al., 2023) used by metaWRAP. Additionally, while metaWRAP runs CheckM repeatedly on each intermediate bin set, Binette uses CheckM2 in a more optimized way, running the most time-consuming component, the DIAMOND part, only once. In comparison, The DAS Tool, which is slightly faster than Binette, also has a good F1 score of 0.771. However, it produces fewer high-quality genomes than Binette (Supplementary Table 5 and Fig. 2).

### “Completion aware” strategy vs. “circular aware” strategy

metagWGS uses the circular contigs as bin sets defined by metaFlye or hifiasm. The Hifi-MAG-pipeline, on the other hand, uses all contigs longer than 500,000 bps as bin sets. If they do not pass the bin quality criteria (> 93% complete), they are then passed on as input, like other contigs, to the binning tools. The previous strategy of the Hifi-MAG-pipeline was based on a “circular aware” strategy and only one binner: MetaBAT 2. The current version uses two binner tools. The novel strategy yields a greater quantity of high-quality bins than the preceding strategy. This is probably due to the fact that the new strategy employs two binning tools and combines their respective results. Indeed, with their new strategy, if a contig with more than 500,000 bps passes the quality criteria (93% of completion), it is considered complete, even if another small contig could complete the MAG.

### Conclusion

metagWGS is the only workflow currently available that provides both the taxonomic affiliation of MAGs, the taxonomic affiliation of all contigs (even those that are not binned) and the functional annotation and taxonomic affiliation of genes.

It is also the only complete workflow for analyzing metagenomic sequencing data in PacBio HiFi. It produces more medium- and high-quality bins than the Pacbio HiFi dedicated binning workflow. Indeed, it uses Binette, which performs better (produces more high-quality bins) than the DAS Tool and the metaWRAP bin_refinement step.

Our ambition is to further develop metagWGS by integrating new tools and strategies, and to keep it up to date with the state of the art.

We plan to investigate co-binning and the tools for using it (semibin2, Vamb - *Nissen et al. 2021*, etc.). Co-binning serves as an alternative to cross-alignment, where reads are aligned on the concatenation of all assemblies. The amount of alignment is the same and the alignment is less parallelized in the case of co-binning, which is not advantageous in the case of execution on a cluster. However, it is the genome that is bigger, so the aligners perform better in the case of co-binning. In addition, the co-binning approach poses problems with the redundancy of concatenated assemblies, although it remains an interesting approach that we would like to test, compare and implement if it is relevant. Co-assembly binning is an excellent alternative to decrease this redundancy but it is not always possible. Indeed, when the samples are numerous, highly diverse and very deeply sequenced, the assemblers run into difficulties and require excessive amounts of resources to co-assemble them.

For better consideration of viruses and small eukaryotes, we also plan to first classify contigs between eukaryotic, prokaryotic and viral contigs and then adapt the tools for structural and functional annotation of genes, select adapted binners, and adapt the tools for taxonomic affiliation and quality assessment of bins/MAGs.

More generally, we plan to continue improving the workflow to make it even faster and less resource-intensive without compromising the quality of the results.

The workflow is available under the GNU GPL license in the following *GitLab* repository: https://forgemia.inra.fr/genotoul-bioinfo/metagwgs. It comes with full documentation, configuration files that you may need to adapt to your infrastructure, and two singularity images to facilitate quick deployment.

## ACKNOWLEDGEMENTS

We would like to thank Marie-Stéphane Trotard for her help on the computing infrastructure. We also thank Maria Bernard for her suggestions that made it possible to improve the workflow and Gail Wagman for proofreading the English.

## SUPPLEMENTARY DATA

**Supplementary Table 1: The main features of seven metagenomic workflows, both functional and technical.**

**Supplementary Table 2: Metrics of assemblies, annotation and reads assigned to genes obtained by metagWGS v2.4.2 from the 11 metagenomic samples sequenced with PacBio Sequel II (HiFi reads).** From left to right: sample name, median read length (after cleaning step), number of HiFi reads in millions, N50 of the assembly, total assembly length, % reads mapped on the assembly, number of contigs, number of CDS annotated, % reads assigned to a gene by featurecount, number of reads assigned to a gene in millions. To obtain these metrics, MetagWGS uses: Quast v5.2.0,minimap2 v2.24 and samtools v1.15 and featureCounts v2.0.3.

**Supplementary Fig. 1: Number of contigs found for each assembly for 11 HiFi public samples from human gut, broken down by length.** The assemblies are made by metagWGS v2.4.2 with hifiasm v0.13-r308, the metrics are calculated with Quast v5.2.0, and the graph itself is generated by MultiQC v.1.14. The samples are represented on the y-axis, and the number of contigs is represented on the x-axis according to their size.

**Supplementary Fig. 2: Cumulative length of sequences for bins of 11 HiFi public samples from the human gut by quality category according to MIMAG (minimum information about a metagenome-assembled genome) standards.** “High-quality” refers to genomes with Completeness > 90% and Contamination < 5%. “Medium-quality” for genomes with Completeness > 50% and Contamination < 10%. “Low-quality” for genomes with Completeness < 50%. “High-contamination refers to genomes with Contamination > 10%. Completeness refers to the proportion of presence of universal single-copy “marker” genes within a genome. Single-copy marker genes present multiple times within a recovered genome is used to estimate potential Contamination. Bins are made by metagWGS v2.4.2, more precisely with Concoct v1.1.0, Metabat v2:2.15, Maxbin v2.2.7, Binette v0.1.5. Bin quality is computed by checkM2 in Binette. The “not-binned” part refers to the cumulative length of assemblies contained in unbinned contigs. The x-axis corresponds to the percentage of the total length of the assembly; the y-axis indicates the samples.

**Supplementary Fig. 3: Quality of dereplicated bins generated from the 11 HiFi public samples from the human gut in terms of completeness and contamination calculated by Checkm2.** The points are colored according to their quality, according to the MIMAG (minimum information about a metagenome-assembled genome) standards. Genomes with the best quality (100% completeness and 0% contamination) are located in the lower right corner of the graph. “High-quality” refers to genomes with Completeness > 90% and Contamination < 5%. “Medium-quality” for genomes with Completeness > 50% and Contamination < 10%. “Low-quality” for genomes with Completeness < 50%. “High-contamination refers to genomes with Contamination > 10%. Completeness refers to the proportion of presence of universal single-copy “marker” genes within a genome. Single-copy marker genes present multiple times within a recovered genome is used to estimate potential Contamination. Bins are made by metagWGS v2.4.2, more precisely with Concoct v1.1.0, Metabat v2:2.15, Maxbin v2.2.7, Binette v0.1.5 and dereplicated with dRep v3.0.0. Bin quality is computed by checkM2 in Binette.

**Supplementary Table 3: Metrics for the 246 MAGs dereplicated to 95% ANI obtained by metagWGS v2.4.2 (with dRep 3.0.0) from the bins for each of the 11 HiFi public samples from the human gut.** From left to right: genome identification and taxonomic affiliation by GTDBtk v2.1.1, completeness and contamination computed by checkM2, genome length, genome N50 and number of reads mapped on contigs making up the MAG, and average depth of alignment of reads on the contigs making up the MAG (by minimap2 v2.24 and samtools v1.15.1).

**Supplementary Table 4: Number of bins produced by the two workflows metagWGS v2.4.2 and Hifi-MAG-pipeline for the 11 HiFi public samples from the human gut.** Number of medium-quality bins for each sample (with more than 50% completeness and less than 10% contamination) by the two workflows and number of high-quality bins produced for each sample with more than 90% completeness and less than 5% contamination by the two workflows. Bins were evaluated by checkM v1.2.2. The highest number is on a yellow square; the lowest on a dark purple square. When the two numbers are the same, the squares are light purple.

**Supplementary Table 5: Comparison of binning results using the simulated mouse gut metagenome data released in preparation for the second round of CAMI II challenges.** To make the comparison, we followed the step in this nature protocol describing the binning benchmark (*Meyer et al., 2021*; *Meyer et al., 2022*). We compared individual binning (Concoct, MaxBin and MetaBAT) and refinement tools (DAS Tool, metaWRAP and Binette). The metrics presented in this table were produced by AMBER v2.0.4. The best result of each metric is indicated in bold.

