## Supplementary figures and images for "metagWGS, a comprehensive workflow to analyze metagenomic data using Illumina or PacBio HiFi reads"

### Supplemental Figure 1

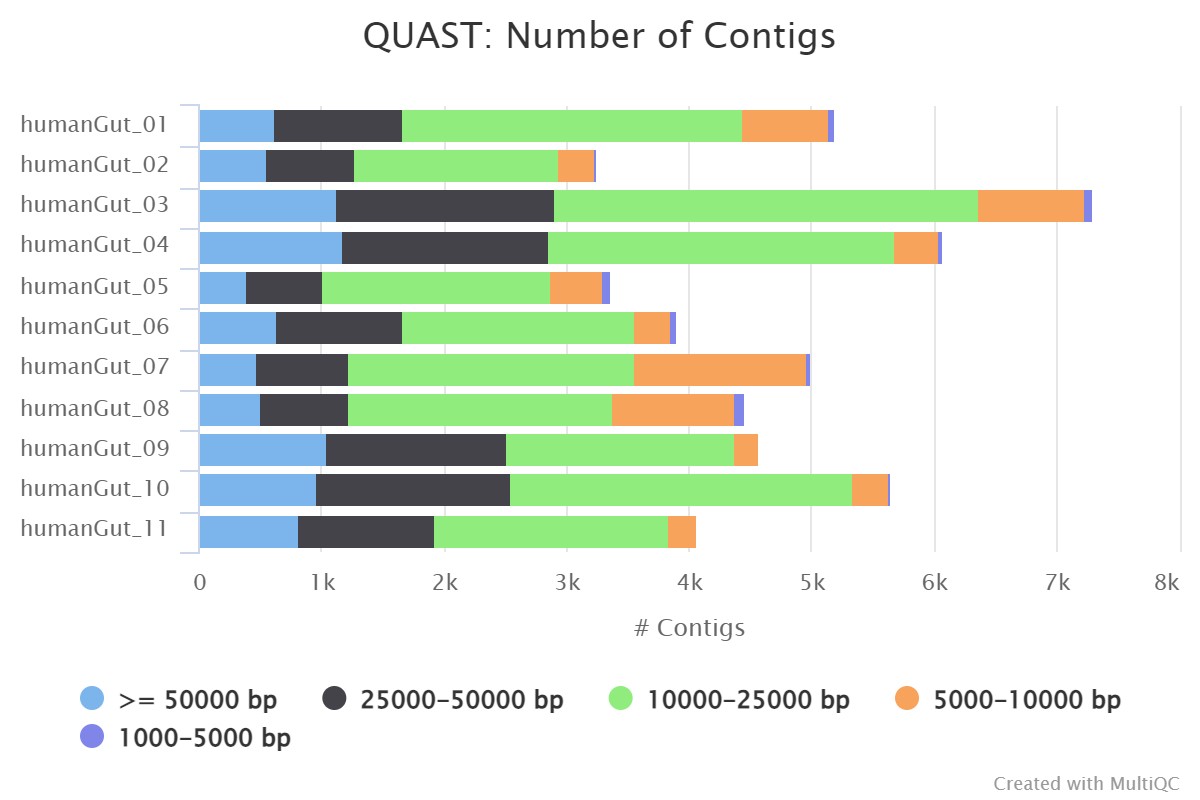

### Supplemental Figure 2

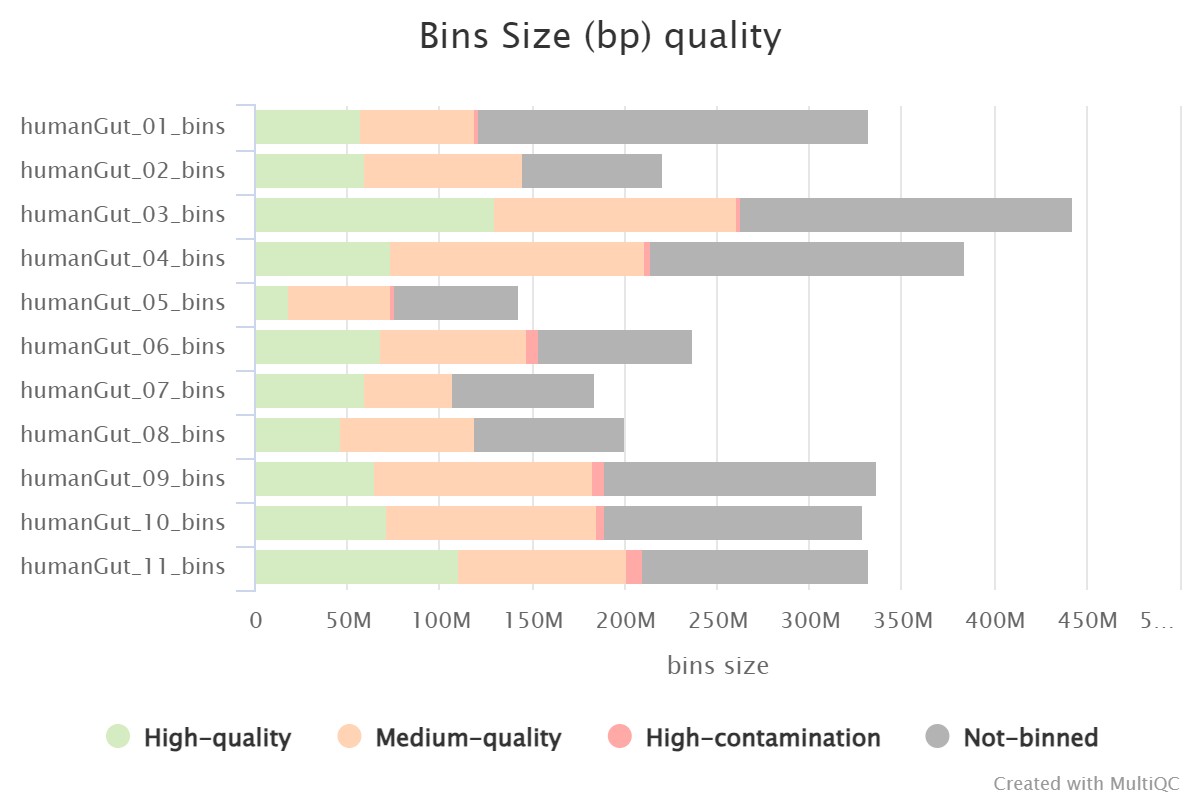

### Supplemental Figure 3

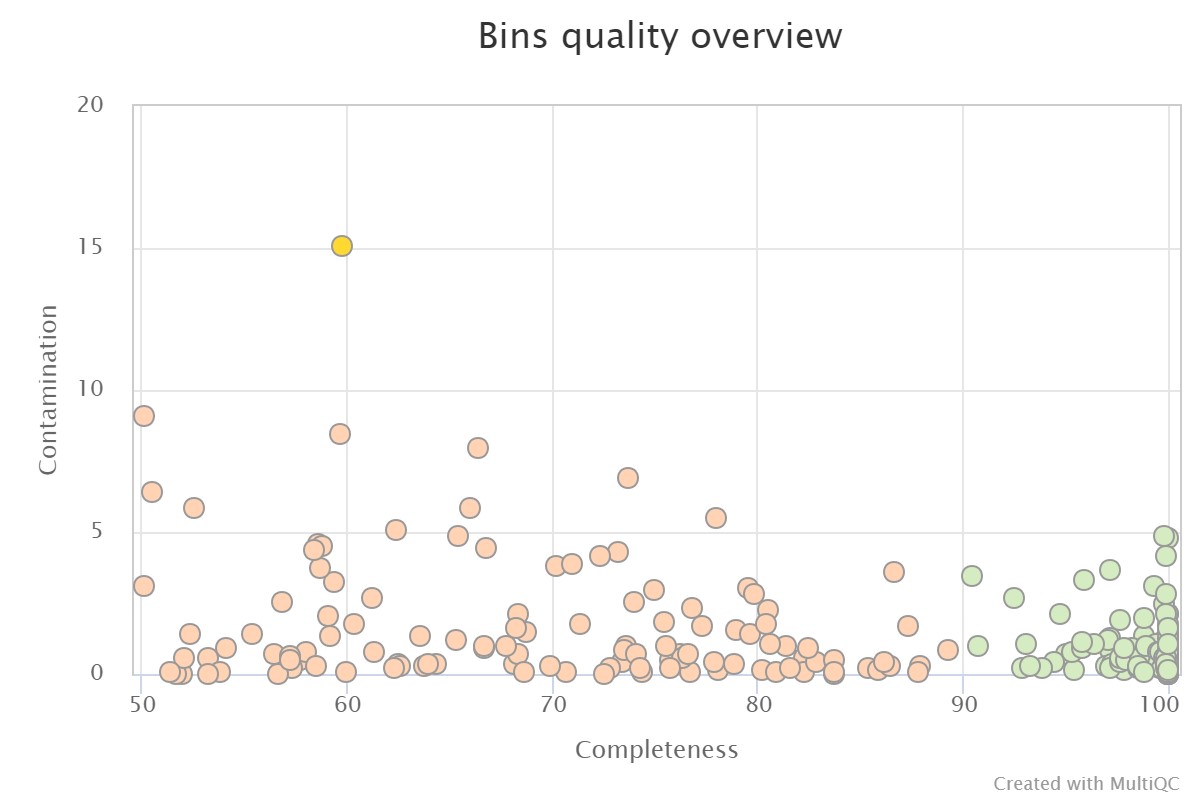
