## Supplemental Table 1 for "metagWGS, a comprehensive workflow to analyze metagenomic data using Illumina or PacBio HiFi reads"

| Workflow | version | repository | website | last update when we did this table | packages / softwares management | pipeline manager and/or languages | Features | ReadQC | Assembly | Structural and functional annotation, taxonomic affiliation | Binning | Reporting | Publication |
| --- | --- | --- | --- | --- | --- | --- | --- | --- | --- | --- | --- | --- | --- |
| MAG | 2.3.2 | https://github.com/nf-core/mag | https://nf-co.re/mag | June 2023 | conda, docker, singularity, podman, shifter, charliecloud | Nextflow, python | Short reads , short and long reads ONT (hybrid mode) | short reads (required):  - fastp (read quality trimming, adapters)  - bowtie2 (host read removal, remove PhiX)  - FastQC (Evaluation)    long reads (optionnal):  - Porechop (Quality trimming, adapters)  - Filtlong (quality filtering)  - NanoLyse (remove Lambda)  - NanoPlot (evaluation)  Taxonomic affiliation of reads  - Kraken2, Centrifuge (taxonomic classification)  - Krona (visualization) | Sample or group-wise assemblies  - Assemblers: SPAdes or MEGAHIT. Hybrid assembler: SPAdesHybrid  - QUAST (Evaluation)  - pyDamage, Freebayes, BCFTools (ancien DNA validation) | ASSEMBLIES:  - Prodigal (protein-coding prediction)  BINS:  - CAT, GTDB-Tk (Taxonomic classification)  - PROKKA (Genome annotation) | - Individual binning tools: MetaBAT2, MaxBin2  Concoct  - Bins quality improving and evaluation: DAS_tool (consolidation binning tool), BUSCO QUAST (evaluation)  checkM and Gunc optionally  - Bins annotation: GTDB-Tk and/or CAT  - abundance estimation  Sample-binning, Group-binning (co-abundance groups defined in config) and all-binning (all samples) | MultiQC | 10.1093/nargab/lqac007 |
| metagWGS | 2.4.2 | https://forgemia.inra.fr/genotoul-bioinfo/metagwgs |  | June 2023 | Singularity | Nextflow, python, bash | Short or long reads (HiFi) (Not hybrid analysis) | - Reads quality trimming and adapters (Cutadapt, Sickle)  - Supress host contaminants: BWA-MEM2 (short reads) or Minimap2 (long reads), + Samtools  - Evaluation: FastQC  - Taxonomic classification of cleaned reads: Kaiju MEM, kronaTools | sample assembly and co-assembly  - Assembly: MetaSPAdes or Megahit (short reads), Hifiasm_meta, metaFlye (long reads)  - Evaluation : metaQUAST  - reads deduplication : bwa-mem2 (short reads) or minimap2 (long-reads) + Samtools  Possibility of giving the assembly as workflow input | - Structural annotation of genes (prodigal) with code genetic alternative (mainly alternative initiation codon : https://www.ncbi.nlm.nih.gov/Taxonomy/Utils/wprintgc.cgi 4 and 11) + tRNAScan + Barrnap  'Functional annotation:  - sample and genes clustering (cd-hit)  - quantifying reads that align the genes (featureCounts)  - Functional annotation of genes, reads quantification per function (eggNOG-mapper)  Taxonomic affiliation:  - Samtools depth, home-made scripts (LCA algorithm) | - Individual binning tools : MetaBAT2, MaxBin2, CONCOCT  - Bins quality improvements and evaluation: bin_refinement module (modified), CheckM2  - Dereplication step (dRep)  - Bins classification : GTDB-Tk  - Bins quantification: BWA-MEM2 (short reads) or Minimap2 (long-reads) + SAMTOOLS  Sample-binning, Group-binning (co-abundance groups defined in config) and all-binning (all samples) | MultiQC |  |
| metawrap | 1.3 | https://github.com/bxlab/metaWRAP |  | août 2020 | conda, docker | Bash, python, R | Short reads | TrimGalore (read trimming and index removal)  bmtagger (remove human reads)  Evaluation: FASTQC | metaSPAdes or MegaHit (assembler)  Filtering step: removes contigs < 1000bp  QUAST : Quality control | Contig blast taxonomy (blobology_module). MEGABLAST, bowtie2, samtools  kraken_module: KRAKEN, KRONA TOOLS (visualization) | Individual binning tools: MaxBin2, CONCOCT, metaBAT2  Bins quality improving and evaluation: bin_refinement module, checkM. Reassemble_bins module, BWA, SPAdes, checkM  Bins quantification: quant_bins module, salmon  Bins classification: classify_bins module, taxator-kt, megablast  Bins annotation: annotate_bins module, PROKKA |  | https://microbiomejournal.biomedcentral.com/articles/10.1186/s40168-018-0541-1 |
| Atlas | 2.17.2 | https://github.com/metagenome-atlas/atlas | https://metagenome-atlas.github.io/ | juillet 2023 | Conda | Snakemake, Python | Short reads , short and long reads (hybrid mode) | Quality control of raw sequence data is performed using BBTools suite:  - clumpify: remove PCR duplicates  - BBduk: remove known adapters, trim and filter reads based on their quality and length. error-correct overlapping paired-end reads  - BBSplit remove contaminating reads using reference sequences (PhiX) | - metaSPAdes or MEGAHIT (short and hybrid assembly)  - contigs abundances (cleaned-reads mapped to assembled contigs) BBmap | - Gene prediction (prodigal)  - Cluster redundant genes (linclust/cd-hit)  - Gene annotation (eggNOG and DRAM)  - Gene quantification by mapping reads to the genes (cds sequences) | '- Individual binning tools: MetaBat2, MaxBIN2, vamb and SemiBin)  - Consolidation binning tool: DAS_tool  - Quality assessment (CheckM)  - Dereplication (dRep)  - Quantification  - Taxonomic annotation : GTDB-tk, GTDB + CheckM  Sample-binning and Group-binning (co-abundance defined in config) |  |  |
| Anvi'o metagenomic workflow | 7.1 | https://github.com/merenlab/anvio/ | https://anvio.org/  https://anvio.org/help/main/workflows/metagenomics/ | October 2021 | Conda | Python, JavaScript, and C  snakemake for the workflow | Short reads | Quality control of metagenomic short reads using illumina-utils, and generating a comprehensive final report for the results of this step  Taxonomical profiling of short reads using krakenuniq. | Individual or combined assembly of quality filtered metagenomic reads using either megahit, metaspades, or idba_ud.  There's a reference mode if reference genomes are known.  Removing the host's reads is possible. | the annotation of your contigs database(s) with functions, HMMs, and taxonomy.  Gene prediction : prodigual  Annotation (HMM against A properly setup local SCG taxonomy database (extracted from GTDB) to quickly estimate the taxonomy of genomes, metagenomes, or bins stored in your contigs-db et ncbi COGs for functions)  quantification by alignment: bowtie2 | All samples can be mapped against all assemblies "all against all".  For automatic binning : You have to option to use several different clustering algorithms, which you’ll specify with the driver parameter: concoct, metabat2, maxbin2, dastool, and binsanity.  The strength of Anvio is its interactive mode that allow manual binning or manual curation of bins |  | https://peerj.com/articles/1319/  https://www.nature.com/articles/s41564-020-00834-3 |
| HiFi-MAG-Pipeline | 2.0.2 | https://github.com/PacificBiosciences/pb-metagenomics-tools | https://github.com/PacificBiosciences/pb-metagenomics-tools/blob/master/docs/Tutorial-HiFi-MAG-Pipeline.md | may 2023 | Conda | Snakemake, Python | Long Reads HiFi | NA | NA | NA | Long contigs >500kb that are > 93% complete by CheckM2) are moved directly to the final MAG set. The others are subjected to binning. Binning algorithms include MetaBat2 and SemiBin2 (using long read settings). The two bin sets are compared and merged using DAS_Tool. Taxonomic assignment: GTDB-Tk. | Various plots |  |
| VEBA | 1.2.0 | https://github.com/jolespin/veba | tutorial : https://github.com/jolespin/veba/blob/main/walkthroughs/end-to-end_metagenomics.md | July 2023 | Docker | Python, genoPype (home-made workflow manager) | Short reads | - Adapter removal and quality trimming. (fastp)  - Pairing of the trimmed reads (BBtools, repair.sh)  - Contamination removal (bowtie2 , with contamination database as input)  - if a k-mer reference database is provided, trimmed or decontamined reads are aligned against the database (BBTools, BBDuk.sh). The read sets are then quantified using SeqKit for accounting purposes (e.g., % contamination or % ribosomal). | - assembles reads (metaSPAdes, SPAdes or rnaSPAdes)  - aligns the reads using to the assembly (Bowtie2)  - Coverage is calculated for contigs via Samtools and genome spatial coverage is provided. Reads from the sorted BAM files are then fed into featureCounts to produce gene-level counts, orthogroup-level counts, MAG-level counts, and SLC-level counts.  - assembly quality control statistics (SeqKit) | - Coverage calculation: align the reads from all provided samples and create bam files (bowtie2, samtools).  - Clusterize genomes (fastANI) and genes from MAGs (OrthoFinder)  - contigs taxonomic annotation (only contigs in bins)  - Genes (from MAGS) functional annotation with various database | - Individual binning tools: MaxBin2, MetaBAT2, CONCOCT  - unbinned contigs are concatenated together to produce a pseudo-coassembly ad then used as input for the binning  - Aggregate binning tool: DAS_tool  - Fast coverage calculations (CoverM)  Binning prokaryote:  - removing eukaryotes at the MAG level (Tiara) : use it to bin eukaryotic genomes with exon-aware gene modeling and lineage-specific quality assessment (metabat2, concoct, coverM, busco, metaEuk for annotation)  - Assess bins quality and remove poor quality bins (CheckM)  Binning eukaryote:  - produce prediction probability for each contig. Contigs from eukaryote MAGs are input into MetaEuk's easy-predict workflow.  - Assess bins quality and remove poor quality bins (CheckM)  Binning viral:  - Potential viral contig are found by VirFinder and verify by checkV, annotation is made by pyrodigal. Unbinned contigs are given as an input at this step (at the eukaryotic binning step)  Assess bins taxonomy :  - GTDB-tk for prokaryotes, checkV for virus and MetaEuk and Busco for eucaryotic genome  No analysis of non-binned contigs | NA | https://bmcbioinformatics.biomedcentral.com/articles/10.1186/s12859-022-04973-8 |
