## Supplemental Table 4 for "metagWGS, a comprehensive workflow to analyze metagenomic data using Illumina or PacBio HiFi reads"

|  | **Medium quality** | | **High quality** | |
| --- | --- | --- | --- | --- |
| **sample** | **metagWGS** | **HiFi-MAGS-pipeline** | **metagWGS** | **HiFi-MAGS-pipeline** |
| humanGut_1 (SRR15489020) | 44 | 49 | 19 | 21 |
| humanGut_2 (SRR15489019) | 51 | 51 | 21 | 19 |
| humanGut_3 (SRR15489018) | 97 | 88 | 47 | 42 |
| humanGut_4 (SRR15489017) | 82 | 84 | 26 | 21 |
| humanGut_5 (SRR15489016) | 27 | 28 | 8 | 5 |
| humanGut_6 (SRR15489015) | 48 | 48 | 24 | 18 |
| humanGut_7 (SRR15489014) | 33 | 34 | 17 | 15 |
| humanGut_8 (SRR15489013) | 43 | 36 | 16 | 15 |
| humanGut_9 (SRR15489011) | 71 | 63 | 25 | 23 |
| humanGut_10 (SRR15489010) | 73 | 75 | 27 | 27 |
| humanGut_11 (SRR15489009) | 68 | 71 | 33 | 34 |
| **Total** | **637** | **627** | **263** | **240** |
